# Growth zone segmentation in the milkweed bug *Oncopeltus fasciatus* sheds light on the evolution of insect segmentation

**DOI:** 10.1101/327106

**Authors:** Tzach Auman, Ariel D. Chipman

## Abstract

One of the best studied developmental processes is the *Drosophila* segmentation cascade. However, this cascade is generally considered to be highly derived and unusual. We present a detailed analysis of the sequential segmentation cascade of the milkweed bug *Oncopletus fasciatus*, as a comparison to *Drosophila*, with the aim of reconstructing the evolution of insect segmentation. We analyzed the expression of 12 genes, representing different phases during segmentation. We reconstruct the spatio-temporal relationships among these genes And their roles and position in the cascade. We conclude that sequential segmentation in the *Oncopeltus* germband includes three phases: Primary pair-rule genes generate segmental gene expression in the anterior growth zone, followed by secondary pair-rule genes, expressed in the transition between the growth zone and the segmented germband. Segment polarity genes are expressed in the segmented germband. This process generates a single-segment periodicity, and does not have a double-segment pattern at any stage.

## Introduction

A defining feature of the arthropod body plan is its segmental organization. The segments - repeating morphological units along the anterior-posterior axis-are formed in a process known as segmentation. The formation of segments occurs very differently in different groups of arthropods. While there is no doubt that segments are homologous among all arthropods, when looking across their full phylogenetic spread, there is relatively little in common in the segmentation process. Nonetheless, from fruit flies to spiders to centipedes, segments are established utilizing a conserved set of transcription factors and signaling pathways, albeit, in different embryonic and cellular environments. Mapping gene-expression patterns during segmentation, in organisms representing key points in the phylogeny and evolution of arthropods, enables the identification of conservation and divergence in the roles of relevant genes, and enables insights into the interplay between them, their functions in segmentation, and the way they have evolved to enable the different observed modes of segmentation.

The segmentation process has been best studied in the fruit fly *Drosophila melanogaster* (1). *Drosophila* segmentation is the text book example of a simple embryological patterning system and is taught in virtually every developmental biology course worldwide. Despite its canonical status in developmental biology, it has been known for over two decades that the *Drosophila* pathway is unusual among arthropods and is highly derived (2). It is therefore extremely interesting to understand how this derived process evolved from the ancestral arthropod mode. Segmentation in *Drosophila* is more or less simultaneous, and is effected through a series of tiered sets of genes, dividing the embryo into smaller and smaller units, culminating in a set of genes expressed in every segment (1, 3). This process and the genes involved therein are usually referred to as the “segmentation cascade”. In contrast, the segmentation process in many arthropods is sequential, with segments being formed one or two at a time, from a posterior growth zone (also known as the “segment addition zone”). Stahi and Chipman (4), traced the evolution of these two modes of segmentation across insects, and showed a complex evolutionary history, including intermediate forms between the two, and cases of parallel gains and losses of both. They suggest that the roots of the *Drosophila* segmentation cascade appear early in evolution, before the radiation of holometabolous insects (those insects with a bi-phasic life cycle punctuated by dramatic metamorphosis). Their interpretation is consistent with an idea originally put forward by Peel (5), according to which there was a gradual transition of control over segmentation from an ancestral posterior cycling mechanism to a gap-gene based simultaneous patterning mode.

Our model organism of choice is the milkweed bug *Oncopeltus fasciatus* (6). *Oncopeltus* is a member of Paraneoptera, which is the sister group to Holometabola. As such, it is ideally situated as an outgroup to the hyper-diverse and widely studied holometabolous insects, and can serve to polarize changes in the segmentation program in a comparison between the two most widely studied insects: *Drosophila* and the flour beetle *Tribolium castaneum*, as well as other holometabolans. Previous work on *Oncopeltus* has also shown that it tends to be fairly conservative, and represents many ancestral characteristics in its developmental program (6). The anterior segments in *Oncopeltus* are patterned simultaneously, through a process that bears many similarities to the *Drosophila* cascade. Posterior segments are patterned sequentially from a growth zone. Our previous work on the *Oncopeltus* growth zone (7) showed that it is divided into two functional domains: a posterior growth zone with high levels of cell proliferation and stable gene expression patterns, and an anterior growth zone with dynamic gene expression patterns and a reduced level of cell proliferation. Comparing this organization to that found in other arthropods, we suggested that it is a general feature of sequentially segmenting arthropods.

Our previous work analyzed only a small number of genes in the segmenting growth zone of *Oncopeltus*. This sample allowed us to demonstrate that segments are formed one at a time, unlike the two-segment periodicity found in both *Drosophila* and *Tribolium* (and convergently in geophilomorph centipedes (8)). Indeed, the ortholog of the *Drosophila* pair-rule gene *evenskipped* (*eve*), famously expressed in a two-segment periodicity in the *Drosophila* blastoderm (9), is expressed in every segment in *Oncopeltus*.

In the current work, we have looked at orthologs of several more genes involved in the *Drosophila* segmentation cascade, including most pair-rule genes and segment-polarity genes (the gap genes have been studied in detail previously (10–13)). We have focused on sequential segmentation during the germband stage, taking advantage of the fact that in species with terminal growth, the anterior-posterior axis serves as a proxy for a time axis, with more anterior regions representing later stages in the process of growth and segmentation. This allows us to identify the temporal sequence of gene activation and to extrapolate to the sequence of developmental events involved in generating segments. Adding RNAi mediated knock-down of some of these genes gives additional information about their function. Placing this in the comparative context described above allows us to discuss some of the key steps in the evolution of the segmentation process in insects.

## Results

### Expression patterns of “segmentation cascade” genes

We followed the expression patterns of 12 genes, mostly orthologues of the pair-rule and segment-polarity genes, during the formation of the abdominal segments of *Oncopeltus fasciatus*. Some of these (*eve*, *Dl*, *cad*, *inv*) have previously been described and will only be mentioned briefly, adding details that have not been previously reported. We present them roughly in the order of their appearance, from posterior to anterior. Note that we use the terms pair-rule genes and segment-polarity genes as convenient shorthand for orthologs of genes that have a pair-rule / segment-polarity role in *Drosophila*, and this does not a-priori imply a similar role in *Oncopeltus*.

#### even-skipped (eve)

The expression pattern of *eve* (7, 14) includes a domain of solid expression in the posterior growth zone, and a striped expression domain in the anterior growth zone (Fig. 1A-A’). The number of *eve* stripes early in abdominal segmentation can be as high as five or six, while by the end of the segmentation process, there is only a single *eve* stripe anterior to the solid expression domain. In some stained embryos, the posterior-most *eve* stripe is in contact with the solid expression domain in its medial portion, giving the impression of a stripe “peeling off” from the solid domain (14).

**Fig 1.**
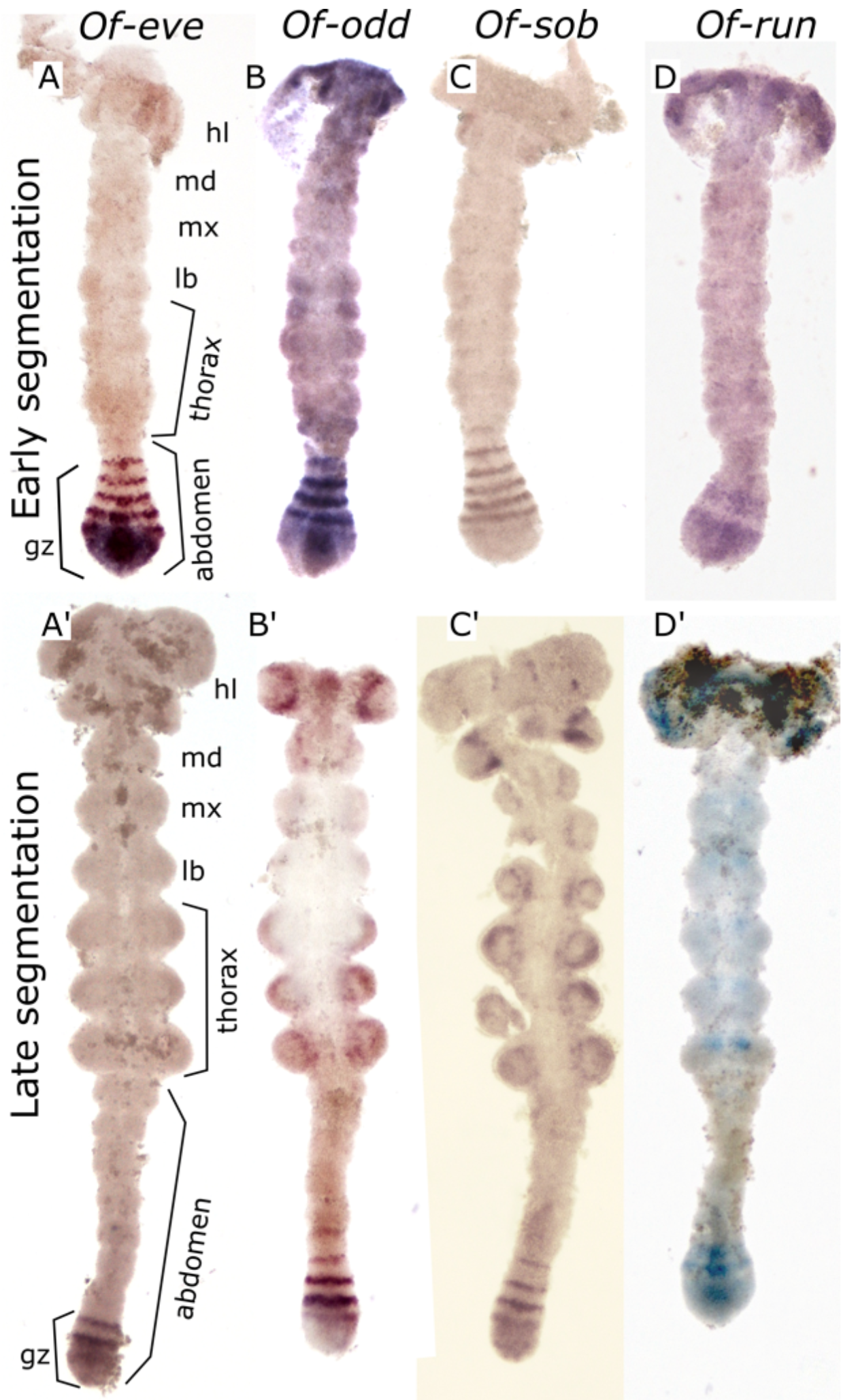
mRNA expression of the pair-rule genes *eve (A-A’)*, *odd (B, B’)*, *sob (C-C’)*, and *run (D-D’)* in embryos at early and late abdominal segmentation. In the early segmenting germband, *eve* (A), *odd* (B) and *sob* (C) all display a similar expression pattern in the anterior GZ composed of 4–6 stripes, corresponding to nascent segments. The main difference between the expression pattern of these genes is most notable in the posterior GZ where *eve* is steadily expressed, whereas *odd* and *sob* show weaker and graduated expression. *run* (D-D’) expression is very different form the other three genes. It is expressed in two broad stripes corresponding to the anterior and posterior GZ, and in patches in the anterior thoracic and gnathal segments. In late germband stages (A’-D’) we see striped pattern of *eve*, *odd* and *sob* maintained, but with a smaller number of stripes. Expression of *run* is decreased to a single broad band in the anterior GZ. In addition, *odd, sob* and *run* are expressed in the limb buds. In all images anterior is to the top. Abbreviations: gz, growth zone; hl, head lobe; md, mandibular segment; mx, maxillary segment; lb, labial segment.

#### *odd-skipped (odd)* and *sister of odd-and bowel (sob*)

The expression patterns of *odd* (fig1B-B’) and its paralog *sob* (fig1C-C’) are nearly identical to each other, and both are remarkably similar to that of *eve*. They also have a solid expression domain in the posterior growth zone, and a striped expression in the anterior growth zone. However, unlike *eve*, the expression of *odd* and *sob* in the posterior growth zone is graded, with highest expression in the anterior margin of the posterior growth zone, tapering off posteriorly, and ending before the posteriormost end of the embryo. The striped expression of *odd* and *sob* extends into the segmented germband slightly more than that of *eve*.

#### runt (run)

Like *odd* and *eve*, *run* (Fig. 1D-D’) is defined as a “primary pair-rule gene” in *Drosophila.* In *Tribolium*, these three genes were found to work together in a pair rule regulatory circuit generating the repeating pattern of the segmentation process (15). The probe for *run* gave very weak signal in our hands, so we could not analyze it at the level of detail we could for other genes. In the *Oncopeltus* germband, *run* does not display a striped expression pattern in the growth zone or in the segmented germband, in contrast with all other “segmentation cascade” genes in this study. It is mainly expressed in two to three broad graduated domains within the growth zone. This pattern is highly dynamic and variable among embryos, but we were unable to correlate this dynamic activity with that of the other genes we have looked at. Expression of *run* can also be seen in the mesodermal cells in the center of the growth zone. In addition to its expression in the growth zone, in late stages of segmentation, *run* is expressed in paired domains in the germband near the ventral midline. The aforementioned four genes are the only pair-rule gene orthologs that are expressed in the posterior growth zone.

#### *odd-paired* (*opa*) and *sloppy-paired* (*slp*)

These two genes are defined as “secondary pair-rule genes” in *Drosophila*. In the *Oncopeltus* anterior growth zone, *opa* is expressed in a striped pattern (Fig. 2A-A’), resembling that of *eve*, *odd* and *sob*. Unlike these genes, *opa* is not expressed in the posterior growth zone at any stage. The number of stripes in the growth zone varies from 2–3 stripes early in the segmentation process to a single stripe at later stages. These stripes are more anteriorly located than the *eve* and *odd/sob* stripes. Expression of *opa* continues into the segmented germband and expression is maintained in narrow stripes in the posterior of each segment throughout the germband stage. The expression of *slp* (Fig. 2B-B’) is similar to that of *opa* with two main differences: the expression stripes are broader in the germband and are found in a more anterior-medial position in each segment. A more subtle distinction is that *slp* has a weak posterior-anterior expression gradient in each stripe, both in the anterior growth zone and in the germband.

**Fig 2.**
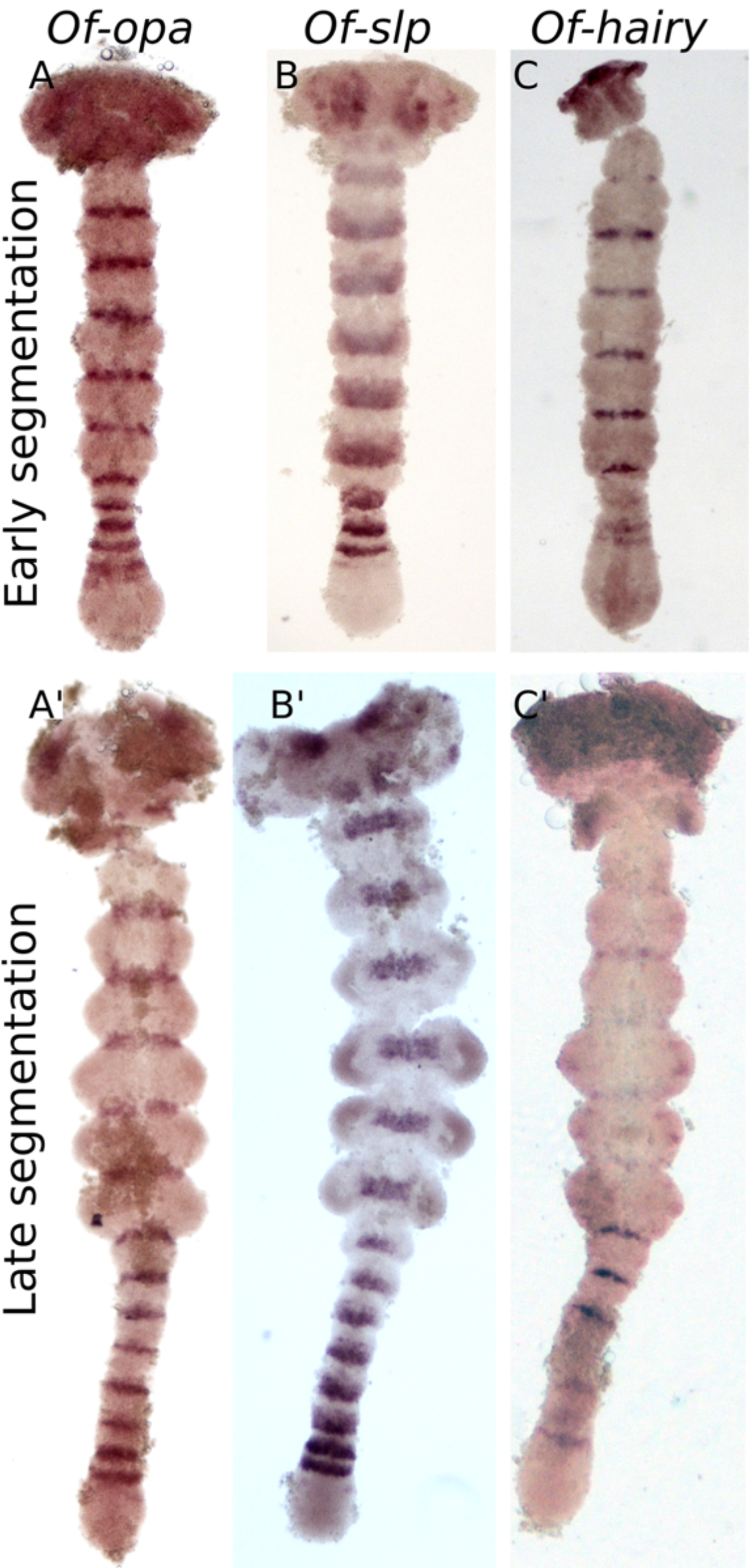
Expression of the pair rule genes *opa* (A-A’), *slp* (B-B’) and *h* (C-C’) in embryos at early and late abdominal segmentation. Throughout development, *opa* (A-A’) is expressed in a narrow band at the border of every segment but is not found in the posterior GZ. *slp* (B, B’) is more broadly and anteriorly expressed in each segment. The earliest, most posterior stripes are thin, and increase in breadth anteriorly. In later stages (B’), it shows diffuse expression in the limb buds. *h* expression (C-C’) is similar to that of *opa* in nascent segments but is weaker in mature segments. In the anterior GZ it is expressed in two stripes at the anterior of the anterior GZ, and more weakly in the posterior GZ. There is also weak punctate expression in the limb buds. In all images anterior is to the top.

#### hairy (h)

Expression of *h* (Fig. 2C-C’) is weakly noticeable in the posterior growth zone of early germband embryos. In the anterior growth zone, it is expressed in two faint stripes, and in a narrow stripe in the posterior of every mature segment. Like *run*, it also shows expression in the mesodermal cells of the growth zone. Segmental expression fades in mature segments later in development.

#### hedgehog (hh)

Known from *Drosophila* as a segment polarity gene, *hh* is expressed not only in every germband segment, but also in stripes in the anterior growth zone, similar to the pair-rule gene orthologs (Fig. 3A-A’). It is visible in the posterior region of the anterior GZ as a wide stripe, not fully resolved and not always clearly separated from the next anterior, better defined stripe. The third *hh* stripe is fully separated from the two prior stripes, as are the more anterior stripes. The segmental stripes are situated in the posterior of each segment. In addition to the striped expression pattern, *hh* is expressed in a single patch at the very posterior of the embryo.

**Fig 3.**
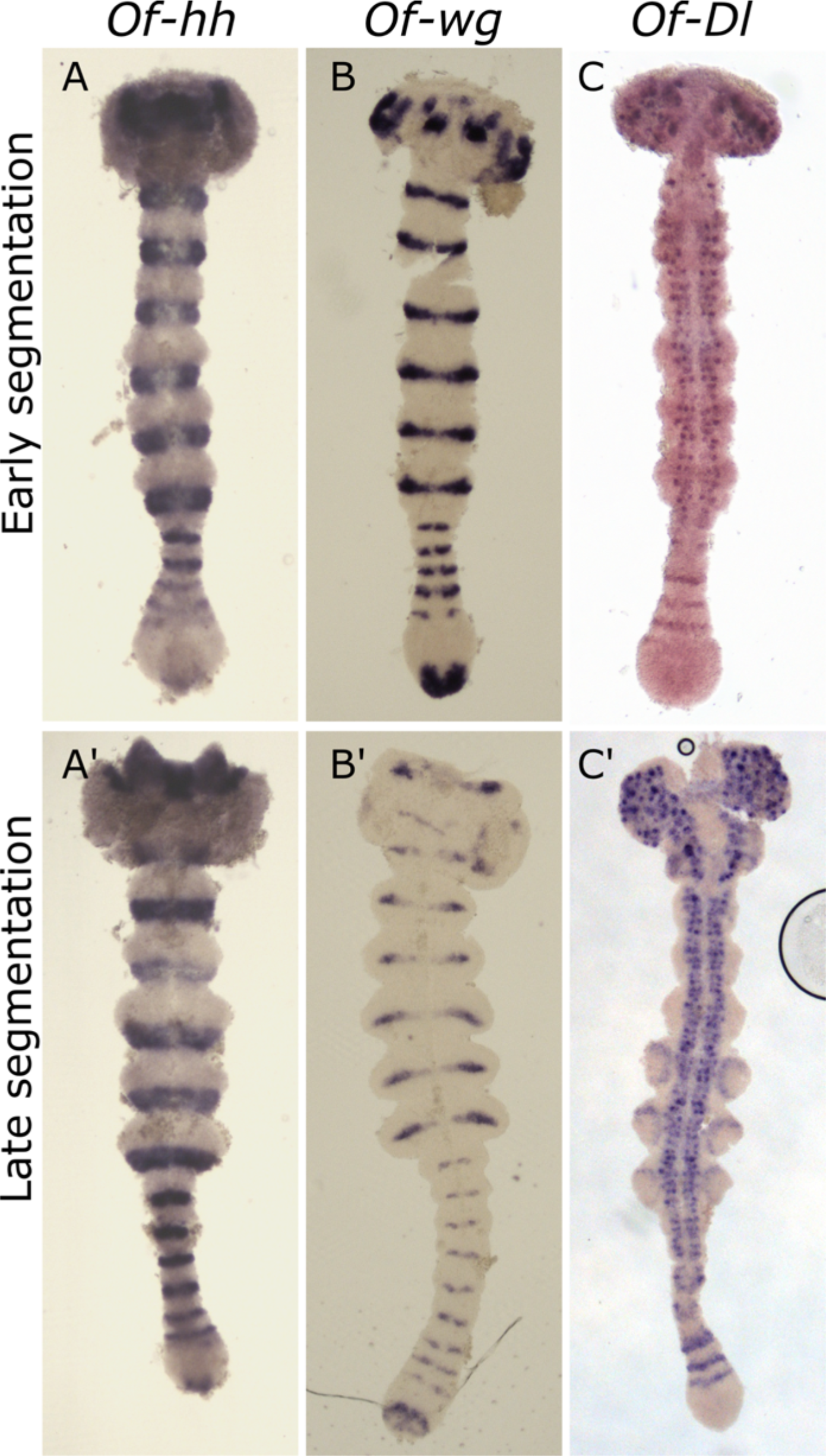
Expression of the segment polarity genes *hh* (A-A’), and *wg* (B-B’), and the Notch ligand *Dl* (C-C)*’*. For the most part *hh* expression corresponds to that of *inv/en*, defining the posterior of each segment. Unlike *inv*/*en*, *hh* is expressed in the anterior GZ, and in a patch at the posterior GZ. *wg* (B, B’) is expressed in the middle of each segment. Like *hh*, it is expressed in a patch in the posterior of the GZ. *Dl* (C-C’) is found to be expressed in a series of stripes in the anterior GZ, and in a punctate pattern in neuronal tissue. In all images anterior is to the top.

#### *wingless (wg)* and *invected (inv)*

We have previously described the expression of *wg* in the *Oncopeltus* blastoderm (4), but not in the germband. Expression of the segment polarity gene *wg* begins in the forming segment, initially as two lateral dots, later expanding and fusing to form a segmental stripe in the middle of each segment (Fig. 3B-B’). The segmental stripes are notable thinner medially. In addition to its segment-polarity pattern, *wg* is strongly expressed in the posterior growth zone. In the early stages of the germband it appears in the posterior pole of the embryo, and as segmentation progresses, it gains a crescent like shape beginning at the medial part of the posterior growth zone, curving anteriorly. At later stages, expression moves slightly anteriorly and gains an M shape. Expression of *inv* (an *engrailed* ortholog) has been described in several previous publications. It is expressed in the posterior of every segment in the germband.

### Relative Expression Domains

To clarify the spatial relationships among the different genes, we carried out a series of double stainings (Fig. 4). These double staining experiments are difficult and unpredictable, and not all combinations of genes were successful. However, we have sufficient pair-wise comparisons to be able to reconstruct the relative position of all of the genes studied (summarized in Fig. 5).

**Fig 4.**
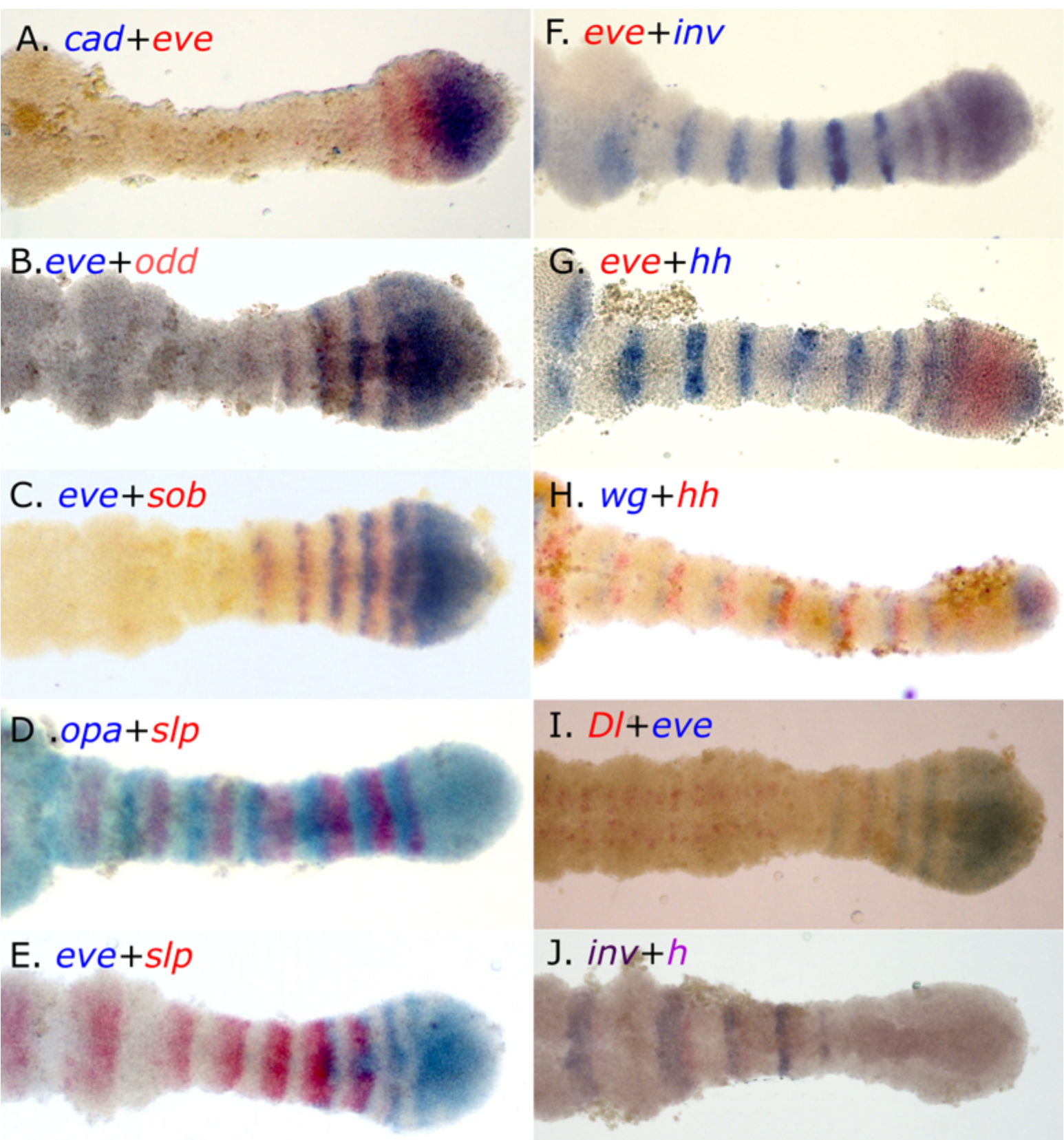
Relative expression domains of different genes in the posterior of the embryo, as illustrated by combinations of double staining. (A) *cad* and *eve* are co expressed in the posterior GZ, with *eve* stripes extending into the anterior GZ. (B) *eve* and *odd*, are shifted relative to each other, with overlapping expression in a narrow area (posterior of *eve* and anterior of *odd*), but with most of the expression separate. (C) The relationship between *eve* and *sob* is identical to that between *eve* and *odd*. (D) *opa* and *slp* are expressed in adjacent domains with no observable overlap. The posterior stripes are complementary and cover the entire anterior GZ. In later, more anterior stripes, as the segment grows, a region without *opa* or *slp* emerges, anterior to *opa* and posterior to *slp.* In later stages, *opa* and *slp* are completely separated. (E) *slp* expression begins just as *eve* expression is fading. In the segments where they are both expressed, *slp* is expressed to the anterior of *eve.* (F) The transition between *eve* and *inv* defines the GZ-germband border. At the transition they are co-expressed in one or two stripes, in which their domains overlap. (G) The first stripes of *hh* expression overlap those of *eve* in the anterior GZ. (H) *hh* is immediately adjacent and posterior to *wg* in segmental stripes beginning in the anterior GZ. This relation is maintained in the posterior GZ, where both are expressed in non-overlapping patches. (I) *Dl* and eve are partially co-expressed in the anterior GZ, with *Dl* extending more anteriorly than *eve*, and eve beginning posteriorly to their overlapping domain. (J) *h* is expressed immediately posteriorly to *inv*, beginning slightly more anteriorly. Anterior is to the left in all images.

**Fig 5.**
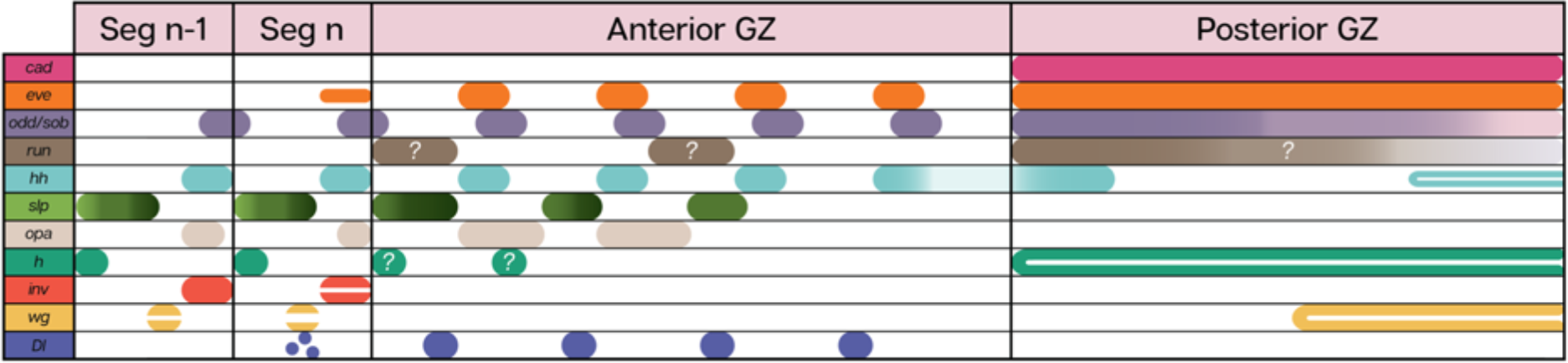
Schematic representation of the relative expression patterns of all the genes discussed, as deduced from the double and single stainings. Question marks indicate cases with ambiguous staining, or where double staining was not possible, preventing us from identifying exact relative expression domains.

Four of the genes we studied are expressed in solid domains throughout the posterior growth zone - *cad*, *eve* and *odd/sob*. We carried out double staining for *cad* and *eve* (Fig. 4A) to see whether they share an anterior border in the posterior growth zone (the border between the anterior and posterior domains of the growth zone). We could detect no difference in the anterior extent of these genes, indicating a single uniform border within the growth zone.

The expression patterns of *eve* and *odd* are similar (Fig. 4B), With full overlap in the posterior growth zone. However, looking at their anterior expression shows that they overlap only partially, with *eve* expression being anterior to that of *odd* in any given stripe. We did not double-stain *odd* and *sob*, however, they both show the same relationship to *eve* (Fig. 4C) suggesting that their expression patterns fully overlap.

The two secondary pair-rule genes, *opa* and *slp* (Fig. 4D) are expressed in complementary patterns in the anterior growth zone, with *slp* forming the posteriormost stripe. As segmentation progresses, in the later stripes of the anterior growth zone, a gap appears anterior to the *opa* stripe and posterior to the *slp* one. In the segmented germband the stripes are fully separated and occupy distinct regions of the nascent segment. Comparing *slp* with *eve* (Fig. 4E) shows that they have a narrow domain in the anteriormost growth zone where both are expressed. Expression of *slp* appears as a faint narrow band in the lateral anterior growth zone, anterior and adjacent to the second *eve* stripe. The second *slp* stripe is already much stronger, but still shows a gap in the midline where *eve* is expressed, indicating that at this stage, these two genes are probably not co-expressed, but are both present in different areas of the same position along the anterior-posterior axis. The third *slp* stripe is completely resolved to the anterior of the final, most anterior *eve* stripe. As visible in the single stainings, *slp* is expressed in a graduated manner, strongly expressed in the posterior of the band, weakening towards the anterior but still with a well-defined anterior border, after which there is a gap where neither *eve* nor *slp* are expressed. The relative expression of *opa* and *slp* suggests that *eve* overlaps *opa* in the anterior of each stripe.

The expression of *eve* and *inv* (Fig. 4F) overlap exactly in the only region where they are co-expressed - the posteriormost segment (as previously shown by Liu and Kaufman (14)). The domains of *eve* and *hh* (Fig. 4G) also overlap, but this overlap extends through the entire anterior growth zone. Thus, we conclude that *inv* and *hh* also overlap. The third segment polarity gene we have looked at, *wg* abuts *hh* and sits anterior to it (Fig. 4H). Thus, the expression of *hh* can be seen as a combination of the posterior expression of *eve* and the anterior expression of *inv*.

Finally, the expression of *Dl* lies anterior to that of *eve* (Fig. 4I), perhaps with a slight overlap. Thus, when it is still expressed in stripes, *Dl* overlaps the expression domain of *slp*. The picture is completed by the anterior expression of *h*, which lies adjacent and posterior to *inv*, but is expressed earlier in any given segment (Fig. 4J).

### Spatial dynamics of the segmentation genes

In order to gain a better understanding of the dynamic pattern of the segmentation genes over time, we have measured the expression levels of three representative genes, *eve, odd* and *hh*, along the anterior-posterior axis (Fig. 6A-C, A’-C’). We summed the pixel intensity for every point along the posterior-anterior axis on photographs of stained embryos. For two of these genes, we followed this up in a large sample of >50 embryos, and plotted summed pixel intensity along the axis over developmental time on a three dimensional graph (Fig. 6D-E. See supplementary methods), including relative developmental age (a value-less order based on germband length), position along the axis (using the posterior boundary of the third thoracic segment as the origin) and normalized expression level. These graphs allow us to follow the development of the expression patterns of these two genes, and by extension, shed some light on the dynamics of the entire process.

**Fig 6.**
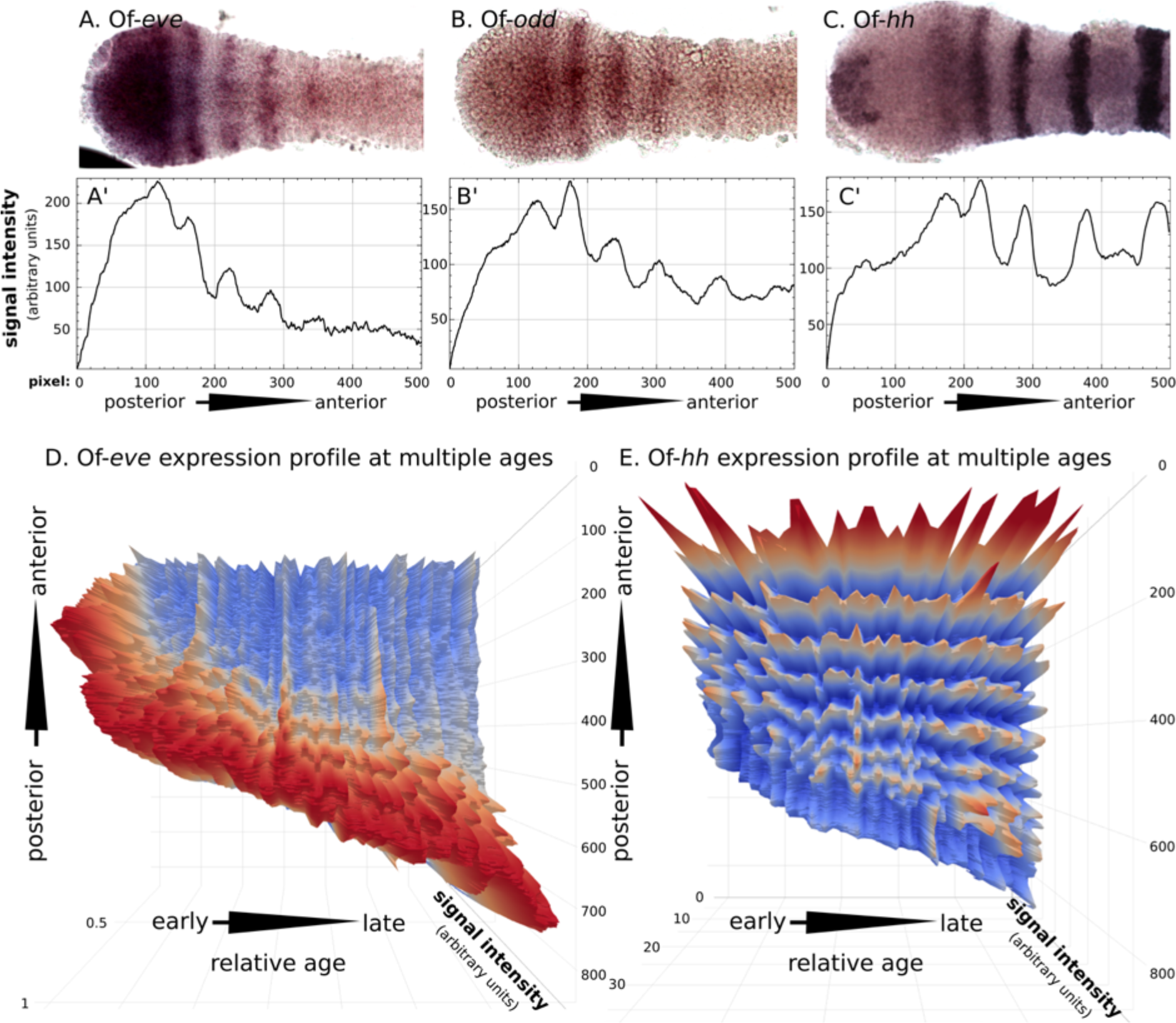
Quantitative analysis of the dynamic expression of *eve*, *odd* and *hh*. (A-C) High magnification images of the growth zone and posterior germband of embryos stained for (A) *eve* (B) *odd* and (C) *hh.* (A’-C’) Gene expression levels in the embryo shown above. For each embryo we drew a rectangle encompassing the entire imaged region and summed the pixel intensities for each point along the x-axis (posterior to anterior). Comparison of the signal intensity highlights the small differences in the expression profile of these genes in the GZ. The main difference seems to be that while *eve* is robustly expressed in the posterior GZ and is strongest in the anterior of the posterior GZ, *odd* is weakly expressed in the posterior GZ, increasing in strength towards the anterior, and peaking only in the first discrete *odd* stripe in the anterior GZ. The main observation regarding the *hh* expression profile is the double-peak between the stripe in the anterior GZ and first stripe of the posterior GZ, which are not completely resolved. Only the third *hh* stripe is completely resolved. (D-E) 3D plots including a sequence of embryos expressing (D) *eve* and (E) *hh*, arranged in sequence by increasing total length of the GZ + abdominal segments. The third thoracic segment (T3) was defined as the anteriormost point of each graph. For *eve* n=71 for *hh* n=53.

Using this visualization, we show that when *eve* stripes first peel off from the posterior growth zone’s solid expression domain, they remain stationary relative to the germband (represented by the third thoracic segment) but shift slightly in position relative to the solid expression domain of *eve* in the posterior growth zone. The stripes of *hh* expression, in contrast, remain in a constant position relative to the third thoracic segment after they are formed.

### RNAi experiments

Following the detailed analysis of gene expression patterns, we went on to examine the function of representative genes in the segmentation process by knocking them down through RNAi. For each gene knocked down, we collected early germband stages and late germband stages of RNAi embryos and stained them for *inv* and *eve*. In addition, we collected prehatching larvae to identify morphological phenotypes.

RNAi experiments have previously been conducted for some of our genes of interest. Knocking down *eve* leads to a truncation of the embryo and a complete loss of all growth zone derived segments (14). Knocking down the segment polarity genes, *inv* and *wg* leads to malformed segmental boundaries, but does not lead to any truncation phenotypes (16). We have knocked down the second important primary pair rule gene *odd*, the secondary pair rule gene *slp*, and the remaining segment polarity gene *hh*.

Early *odd-RNAi* germband embryos (Fig. 7B) exhibit a reduction in the distance between the anterior gnathal and thoracic segments, and a much broader expression of *inv*. The maxilla and labium are closer together with cells between them expressing *inv* ectopically. T1 and T2 are also wider and closer, and are fused in the midline. T3 seems to be normal, and so do the abdominal segments present at this stage. The growth zone exhibits no visible abnormalities. In later, fully segmented germband embryos (Fig. 7B’), we see that this phenotype has progressed to limb fusion: T1 and T2 are fused. T3 remains mostly separated. In the abdominal segments we see a similar effect of segment fusion and occasional ectopic expression of *inv* between segmental stripes. Notably, we see no evidence of segment deletion. The number and location of the segments is normal, but with defective segment boundaries. This is true both for the blastoderm derived anterior segments and for the growth zone derived posterior segments. The larval *odd* RNAi phenotypes are remarkably uniform, and nearly all show the same findings: segments are formed with irregular boundaries, appendages are fused and we see deformation in the head, mainly indicating abnormal midline closure.

**Fig 7.**
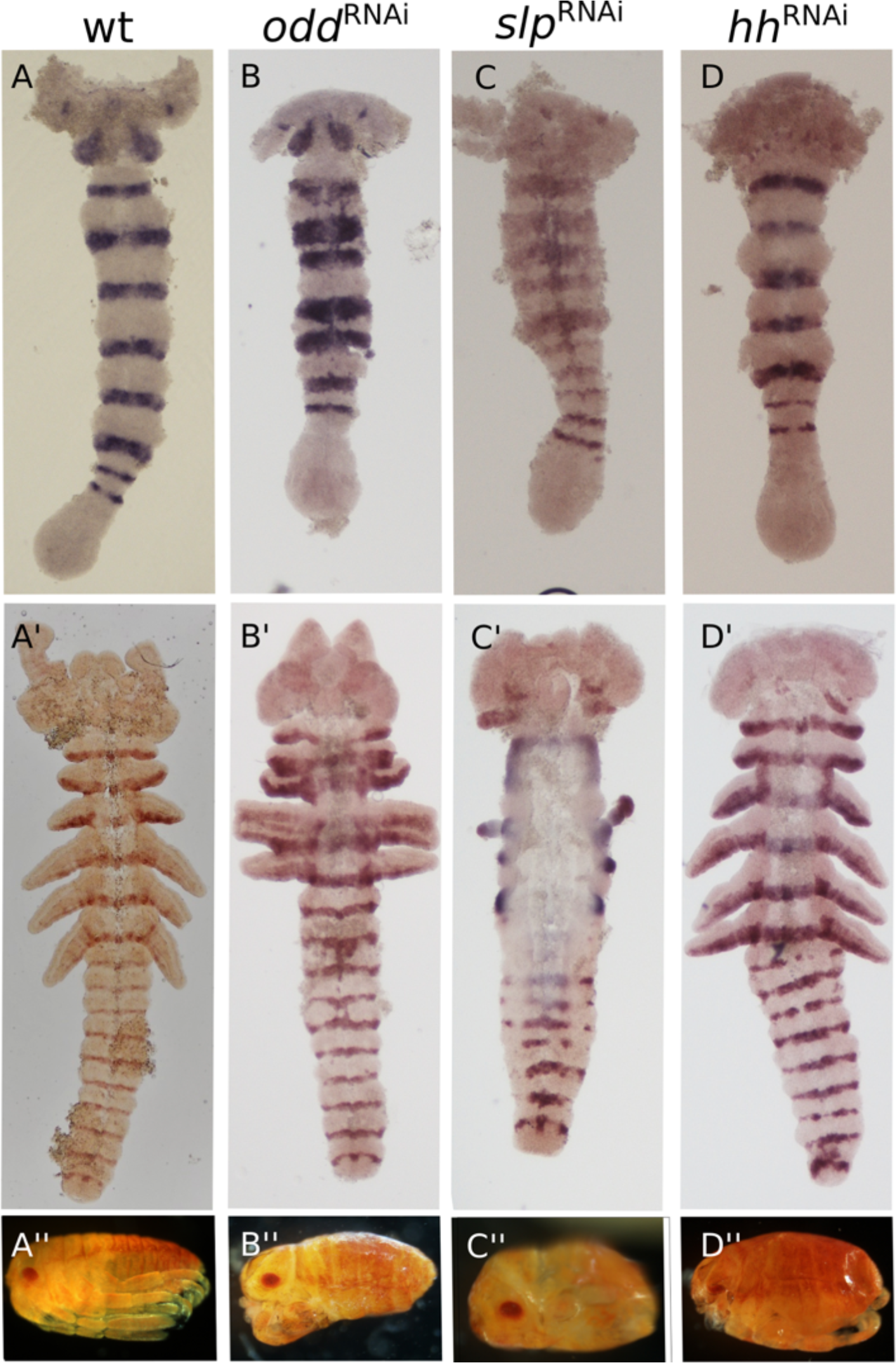
Segmental phenotypes following knock-down of *odd* (B-B”), *slp* (C-C”) and *hh* (D-D”), in early and late germband embryos stained for *inv* and in hatchlings. (A-A’’) Wildtype embryos and hatchling. (B) In the early germband embryo *odd* RNAi embryos mainly display widening of *inv* expression in the thoracic segments, and fusion of segments in the embryonic midline. (B’) In later stages, appendages are fused, and the borders of some abdominal segments are also ill-defined, sporadically fused or narrowed. In both embryonic stages slight ectopic expression of Of-*inv* is seen in single cells. (B’’) In the *odd*-RNAi hatchling this phenotype causes compression of the thorax and truncated limbs. (C) in *slp*-RNAi embryos thoracic *inv* expression is broader in the early germband embryo, and abnormally expressed in the midline. (C’) The later *slp*-RNAi embryo displays severe truncation of all appendages, with only limb buds of T1 and T3 remaining. In addition, we see malformation of the abdominal segment boundaries, where gaps in *inv* expression can be seen. The *slp*-RNAi embryo is also wider than WT embryos and has an apparent breakdown of midline tissues. (C”) The *slp*-RNAi hatchlings are compressed with almost no segmental boundaries, and holes appear in the lateral parts of the embryo, where the limbs are missing. (D) Early hh-RNAi embryos seem to be almost completely normal, only displaying some minor head aberration. (D’) Aberrations of the head are also seen in the late germband embryos which seems to lack some folds and finer details of the head structure. Abdominal segment borders are also affected, containing gaps and ectopic expression of *inv* in sporadic cells. (D”) In hatchlings, the head is greatly reduced and malformed. Segmental borders can be seen, but they are disrupted. Limbs develop normally.

Similarly, *slp* RNAi embryos retain the normal number of segments (Fig. 7C’), but these are misshapen. Already in the early germband embryo (Fig. 7C) the embryos are noticeable wider, and we can see an expansion and ectopic expression of *inv* in the midline of the blastoderm derived segments. At later stages, the ventral (medial) region is much wider and thinner, with unusual excretions obscuring the cells (possibly indicating apoptosis). Expression of *inv* is lost from the ventral portions of these segments (Fig. 7C’). The first abdominal *inv* stripes are expressed relatively normally, and only in the later embryo (Fig. 7C’) do we see that the segment border is malformed in the mediolateral aspect of the abdomen, lacking normal expression of *inv*. The most striking outcome of *slp* RNAi is the loss of thoracic appendages. Some embryos are completely devoid of appendages, while some maintain residual stumps of limbs T1 and T2. This is also seen in the hatchlings (Fig. 7C”), where instead of limbs we find actual holes in the cuticle. We repeated the RNAi experiment with a second fragment. The results were similar, but the phenotypes were generally weaker (Supplementary Fig. 2). Interestingly, both in germband embryos and in hatchlings, we see that the T2 limb is lost before the other limbs.

RNAi for *hh* (Fig. 7D-D”) gave similar results to those previously reported for the other segment polarity genes (16). All segments are present, and germband embryos look almost normal. However, larvae are compressed and show disrupted segmental boundaries. Malformations of the head are seen both in the germband embryos and in the hatchling.

We looked at the expression of *eve* in RNAi embryos for all three genes. In all cases, the expression of *eve* in the growth zone is almost indistinguishable from wildtype expression (Supplementary Fig. 1). However, in *hh-*RNAi embryos, we see ectopic expression of *eve* in a stripe in the head region, and in *odd*-RNAi embryos we see ectopic expression in the midline of the germband.

## Discussion

### *The segmentation “cascade” of* Oncopeltus

Our results, coupled with our previous analyses of segmentation in *Oncopeltus* (4, 7), allow us to reconstruct the series of molecular events involved in defining segments from the growth zone (Fig. 5). The first event in the process is the separation of an *eve* stripe from the posterior growth zone and a limited movement of *eve* expression across cells at the posterior margin of the anterior growth zone. We suggest that this activates the entire downstream sequence of expression patterns. However, we still do not know what generates the repeating process (or oscillator) that causes *eve* stripes to peel off and move anteriorly. The two possible candidates (17) are Delta-Notch signaling, as found in centipedes, spiders and branchiopod crustaceans (18–21), or a pair-rule gene circuit, as found in *Tribolium* (15). Neither one of these candidates is fully consistent with our data. *Dl* is not expressed at the right time and place to be upstream of *eve*, and knocking it down does not disrupt the early segmentation process, but rather the later stages of segmental boundary formation (4). A gene circuit as in *Tribolium* is a possibility, but not with the exact same interactions, since knocking down *odd* expression does not affect *eve* expression in the growth zone, and because *odd* and *eve* expression domains overlap, making a repressive interaction between them unlikely. We also cannot draw clear conclusions about the possible role of *run* in such a circuit due to the poor quality of our *run* staining. Following striped expression of *eve*, several other genes are expressed in a similar domain, including at least *odd, sob* and *hh*. Based on their spatial relationships we suggest that these genes are activated by *eve* (either directly or through a close intermediary), but we cannot test this functionally since knocking down *eve* leads to a complete truncation of the growth zone and sequential segmentation does not take place.

The next phase occurs in the anterior part of the anterior growth zone. Expression of a series of secondary pair-rule gene orthologs is activated, including at least *opa* and *slp*. We suggest that *opa* and *slp* are repressing each other, as there is no overlap in their expression domains at any point. Expression of *hh* is maintained at this stage. Slightly anterior to where the secondary pair-rule genes are activated, expression of *eve, odd* and *sob* is switched off and the segment polarity genes *inv* and *wg* start to be expressed in stripes. Interestingly, *h*, which is a pair-rule gene in *Drosophila* and in *Tribolium* (22), is expressed segmentally late in the cascade, anterior to the expression of *wg* and *inv*, although it has an earlier non-segmental mesodermal expression. The point of activation of *inv* was previously defined as the border between the growth zone and the segmented germband (7). Genes that are expressed in stripes at this border (the segment polarity genes and the secondary pair-rule genes) remain active throughout the germband stage and maintain their striped expression in each segment as the segments continue to mature.

Thus, segmentation from the growth zone occurs through three phases indicated by the expression of orthologs of primary pair-rule genes, secondary pair-rule genes and segment-polarity genes. The boundary between these phases is not sharp, and several genes are active across phases (e.g. *hh*). We have not looked at gap genes in the current work, since this group of genes and its role in segmentation has been studied previously (10–13). In the *Oncopeltus* blastoderm, gap genes have a regulatory role in forming specific segments, as they do in *Drosophila* (4). Existing data do not support a role for gap genes in the sequential segmentation cascade in *Oncopeltus*, since they are not expressed in the growth zone, but only in nascent segments.

### Changes in the growth zone

We have previously documented the changes in size of the growth zone throughout the segmentation process (7). Here we expand on these results by documenting the expression levels of two genes holding central positions in the segmentation cascade, relative to the dynamic growth zone (Fig. 6D-E). Using the posteriormost blastoderm – derived segment – the T3 segment-as a fixed reference point, we show that the growth zone moves posteriorly as the germband elongates, and that nascent segments remain stationary at the point where they were first determined in the anterior growth zone. This leads us to the surprising conclusion that the early stripes of pair-rule gene expression already commit the cells where they are expressed to their future segmental identity. Cells expressing *eve* in the anterior growth zone remain in the same position as they go through the segmentation cascade, ultimately expressing *inv* as the anterior growth zone contracts latero-medially to give rise to a new segment of the germband.

In many species that have been studied, there is a phase wherein there is a wave of cyclical gene expression traveling across cells (8, 23, 24). Our results are not consistent with a longdistance traveling wave in *Oncopeltus*. If there is a movement of expression across cells, it is only at the very early stage of *eve* expression, where a new stripe peels off of the posterior growth zone.

Our RNAi experiments raise an interesting contrast with mutant phenotypes of orthologous genes in *Drosophila*. None of our experiments result in the loss of specific segments or segmental domains. They all exhibit different levels of disruption of the segmental borders or segmental structure. This strengthens our assertion that segmental commitment occurs very early relative to that known from *Drosophila*. Indeed, knocking down *eve*, which we identify as the earliest gene in the cascade, leads to a complete loss of all growth zone derived segments. In addition, the weak RNAi phenotypes indicate a tightly integrated gene regulatory network, with a high level of redundancy.

### Conservation of the segment polarity network

The segment polarity network is generally considered to be the most conserved part of the segmentation process in arthropods (25, 26). This seems to hold for *Oncopeltus*. The expression border between *wg* and *inv*/*en* defines the parasegment boundary in *Drosophila* (27). We find the relative expression pattern of these genes is conserved in *Oncopeltus.* The position of *inv* relative to *wg* and *hh* in the germband segments (Fig. 5) is deduced from the relative expression of *eve* and *inv* and by the co-expression of *eve* and *hh* in the anterior growth zone (Fig. 4G), thus confirming that *eve/inv*, and *hh* are expressed in the same part of the nascent segmental, though not at the same time. Expression of *wg* is adjacent and anterior to these, as in *Drosophila*.

The behavior of *hh* is somewhat different from the two other segment polarity genes we studied, in that its expression begins much earlier in the cascade, concomitantly with the early expression of *eve*. This is similar to the early expression of *hh* reported from the scorpion *Euscorpius* (28). The early expression of *hh* is an interesting indication of the processes of segment maturation. Given its position at the posterior border of the segment, the expression of *hh* indicates that segment polarity and segment boundaries, usually perceived as late milestones in segment maturation, are actually established very early on, almost immediately as the tissue enters the anterior growth zone. Furthermore, when examining the progression of the expression pattern of *hh* (Fig. 6E), we note that the posterior (early) *hh* stripe is broad, not fully resolved and not separated from the second *hh* stripe. Only the third or fourth stripe of *hh*, which coincides with the beginning of the expression of *en*/*inv* and *wg*, is completely resolved and with sharp borders. Similarly, in *Drosophila*, *hh* is initially broadly expressed within the parasegmental unit, and is later refined to a narrower region. In addition to this, as demonstrated by double stainings, *hh* is also expressed in the posterior growth zone, surrounded by a crescent of *wg* expression. The relative expression of *hh* and *wg* in the posterior growth zone is the same as that later observed in the segment border.

### Relative rate of segmentation

In previous work (7) we analyzed the dynamics of sequential segmentation from the growth zone, and showed that the rate of segment generation, using *inv* expression as a proxy, is not significantly different from linear throughout the process. In the present work we were able to look at different phases of the segmentation process. Looking at the number of stripes of *eve* and other genes expressed in the anterior growth zone, we see that this number varies from 4-5 stripes in early stages to only 1 towards the end. We suggest that this indicates different processes that are not temporally linked. The first phase of segment determination, indicated by *eve* expression, occurs very rapidly, creating a “backlog” of segments waiting to go through the next phases, indicated by the expression of secondary pair-rule genes and segment polarity genes. Thus, by the time the first nascent abdominal segment starts expressing *inv* there are already 4–5 subsequent segments expressing *eve*. As segmentation progresses, primary determination slows down and final determination catches up, so that there is only one segment expressing *eve*.

### Evolution of the segmentation cascade

Comparing our findings to what is known from better-studied experimental systems (most notably *Drosophila* and *Tribolium*) allows us to identify key aspects of the process that are broadly conserved at different phylogenetic scales and to reconstruct some of the evolutionary changes that have taken place in the evolution of the segmentation cascade, both within insects, and in arthropods more broadly.

The transcription factors commonly known as pair-rule genes hold a key early role that is conserved in the segmentation cascade of all arthropods studied to date. Although the gene studied most widely has been *eve*, orthologs of other members of this group interact with *eve* in many cases. These genes provide the first reiterated output that sets the path for segment determination. The signal driving pair-rule gene expression is variable and ranges from simultaneous activation in many segments as in *Drosophila* and in the *Oncopeltus* blastoderm (4), through Notch-signaling as in centipedes (20), to an endogenous pair-rule gene circuit as in *Tribolium* (15). Nonetheless, the centrality of pair-rule genes in subsequent stages is conserved. In the case of *Oncopeltus* sequential segmentation, *eve* is very high in the cascade and is most likely upstream of all other segmentation genes.

Within pair-rule genes, a distinction between primary and secondary pair-rule genes is also broadly conserved, although the precise distinction of which genes fall into which category varies among taxa (29). The primary pair-rule genes are active together at early phases of the cascade, while secondary pair-rule genes are active later, and are co-expressed with segment polarity genes further down the cascade.

### The evolution of double-segment patterning

While we have been using the moniker “pair-rule genes”, in reality the two-segment periodicity of these genes is probably taxonomically restricted to holometabolous insects. A two-segment periodicity is also found in geophilomorph centipedes (8), but this is likely to be independently evolved, since there is no evidence for such a periodicity in lithobiomorph centipedes (30), in chelicerates or in crustaceans. Double-segment patterning is common to most holometabolous insects, and we have previously suggested that it appeared at the base of Holometabola (4). Within hemimetabolous insects, the only evidence for a pair-rule periodicity is in the cricket *Gryllus bimaculatus* (31) where some of the *eve* stripes exhibit stripe splitting, suggesting an intermediate step on the way to full two-segment periodicity patterning. Regardless of whether the situation in *Gryllus* represents a novelty for crickets and relatives, or whether it is indicative of an earlier appearance of pair-rule periodicity, the single segment generation mode of *Oncopeltus* is probably representative of the situation from which pair-rule segmentation evolved.

The transition between single-segment patterning and double-segment patterning is significant. In geophilomorph centipedes this transition was probably accompanied by a doubling of segment number (32, 33). In insects, there is no change in segment number, suggesting a very different mechanism. Double-segment patterning in insects requires each expression stripe of the pair-rule genes to translate to half a segment at later stages. Taking the *Oncopeltus* cascade as a hypothetical starting point, we can try to uncover the roots of the transition. The primary pair-rule genes are expressed in almost overlapping domains. However, the secondary pair-rule genes *opa* and *slp* are expressed in mutually exclusive domains, similar to those seen in double-segment patterning. We suggest that this mutually exclusive pattern within a single segment was elaborated to pattern consecutive segments, a process which also included a shift of the primary pair-rule gene expression domains to create non-overlapping sets of odd and even segment genes. Intriguingly, *Oncopeltus* does have one gene, encoding the nuclear receptor E75A, that is expressed in a double segment periodicity and shows a pair-rule phenotype upon being knocked down (34). While we do not know how this gene fits into the segmentation cascade in *Oncopeltus*, it may indicate an early stage of segment identity definition, which was part of the basis for the evolution of a double-segment periodicity in patterning the segments.

The evolution of the double-segment patterning mode in holometabolous insects apparently involved several more fundamental differences in the way segments are patterned, relative to the putative ancestral mode seen in *Oncopeltus*. The definition of the segmental unit is much earlier in the *Oncopeltus* cascade, and is already manifested at the level of the primary pair-rule genes. The subsequent cascade refines the borders and possibly defines domains within the segment. Thus, when any gene in the cascade is knocked down (with the exception of *eve*), we see malformations in the segments and in their borders, but no loss of segments. Conversely, in holometabolous insects, knocking down genes higher than the segment-polarity level leads to loss of specific segments (1, 35–37).

### The evolution of simultaneous segmentation

A key characteristic of the *Drosophila* segmentation cascade is the fact that segments are patterned simultaneously, as opposed to sequentially in the ancestral mode (often referred to as long germ vs. short germ development (38), respectively, but see Stahi and Chipman (4)). The transition between these two modes is not well understood. Two recent papers suggest, based on computational considerations, that the transition is actually fairly simple, and requires only minor changes in relative timing of the inputs to the cascade (39, 40). This is consistent with our observation that many aspects of the sequential segmentation cascade in *Oncopletus* are similar to the simultaneous cascade of *Drosophila*. However, both computational models assume input by gap genes, whereas we have no direct evidence of gap-gene input into the sequential cascade.

### Concluding remarks

We have presented a detailed analysis of the genetic events involved in sequential segmentation in *Oncopeltus*. Coupled with our previous analysis of dynamic morphological events in this species, we provide a reference point for comparison with better studied species, to allow a reconstruction of the evolution of the segmentation process in insects. There is a remarkable degree of similarity among the studied insects at the level of molecular players and the general structure of the cascade, despite minor differences in the detail of network structure and significant differences in the cellular and morphological setting. Given the phylogenetic position of *Oncopeltus* relative to *Tribolium* and *Drosophila*, our analysis provides important insights into the evolution of one of the best studied developmental process-the generation of the segmented *Drosophila* blastoderm – and sheds light on how developmental networks evolve.

## Materials and methods

*Oncopeltus* husbandry, embryo collection, fixation and in situ staining were all performed as described in previous work (10) with the exception of the substrate for AP reaction in the last stage of the staining. For the double stainings, this reaction was done using Vector Labs’ vector blue and vector red substrates.

RNAi experiments were also carried out as previously described (10): dsRNA of the gene of interest was injected to the abdomen of virgin females. Embryos were collected and fixed at the required age, with a few embryos left to fully develop in order to assess the potency and effect of the treatment on the hatchling.

For full clone sequences and dsRNA used for knock down, see Supplementary Methods.

## Acknowledgements

We thank Oren Lev and Netta Kasher for help in preparing figures 5 and 6. This work was funded by a grant from the NSF / BSF IOS program: BSF 2012763, and by an Israel Science Foundation grant #120/16.

